# One-step generation of monoclonal B cell receptor mice capable of class switch recombination and somatic hypermutation

**DOI:** 10.1101/218438

**Authors:** Johanne T. Jacobsen, Luka Mesin, Styliani Markoulaki, Cecília B. Cavazzoni, Djenet Bousbaine, Rudolf Jaenisch, Gabriel D. Victora

**Affiliations:** Laboratory of Lymphocyte Dynamics, The Rockefeller University, New York, NY, US; Center for Immune Regulation, Oslo University Hospital, University of Oslo, Oslo, Norway; Whithehead Institute for Biomedical Research, Cambridge, MA, USA; Carlos Chagas Filho Biophysics Institute, Federal University of Rio de Janeiro, Rio de Janeiro, Brazil; Boston Children’s Hospital, Harvard Medical School, Boston, MA, USA

## Abstract

We developed a method for rapid generation of B cell receptor (BCR) monoclonal mice expressing pre-rearranged *Igh* and *Igk* chains monoallelically from the *Igh* locus by CRISPR/Cas9 injection into fertilized oocytes. B cells from these mice undergo somatic hypermutation (SHM), class switch recombination (CSR), and affinity-based selection in germinal centers. This method combines the practicality of BCR transgenes with the ability to study Ig SHM, CSR and affinity maturation.

## INTRODUCTION

Genetically modified mice expressing predefined monoclonal B cell receptor (BCR) repertoires are essential tools in immunological research. The first monoclonal BCR mice were made by injection of plasmids encoding heavy and light immunoglobulin (Ig) chains that integrated together at random sites in the genome^1^. These mice have greatly advanced our understanding of aspects of immune regulation such as allelic exclusion of antibody V region genes ^2-5^ and B cell tolerance to neo-self-antigens ^6,7^ or true self-antigens^8-10^. Although mice can be generated relatively rapidly using this strategy, a major shortcoming is that, given that the transgenic BCR is expressed from a non-native locus, B cells from these mice cannot undergo class switch recombination (CSR), somatic hypermutation (SHM) or affinity maturation. Therefore BCR transgenic mice cannot be use to study a large fraction of the phenomena of interest to B cell immunologists. To circumvent this issue, a second generation of mice were created in which pre-reassembled V_H_ and/or V_L_ regions are inserted into their native loci by homologous recombination^11,12^. These mice are capable of SHM and CSR, and thus allow a wider range of phenomena to be studied. However, traditional knock-in technology relies on complex genetic manipulation of embryonic stem (ES) cells, and, if fully monoclonal B cells are to be achieved, on the generation of two separate mouse strains (one for the Ig heavy chain (IgH) and one for the Ig lambda/kappa light chain). This approach is costly and labor intensive, in addition to requiring more complex breeding strategies in order to maintain both Ig chains together once the strains are generated.

Recently, site specific CRISPR-Cas9 nuclease has been shown to efficiently induce double-stranded breaks in DNA in fertilized oocytes^13^, enabling homology-directed incorporation of transgenes directly at this stage. We took advantage of this technology to target a bicistronic allele encoding both the light and the heavy Ig chains to the endogenous *Igh* locus. Thus, in a single step, we were able to generate monoallelic BCR monoclonal mice capable of CSR, SHM and affinity maturation, in the same timeframe required for untargeted BCR transgenics.

## RESULTS AND DISCUSSION

We began by determining which single guide (sg)RNAs were optimal for generating double-stranded breaks at the 5’ and 3’ of a ~2.3 Kbp region spanning the four J segments of the *Igh* locus (Fig 1a,b). Cutting efficiency was assayed for several sgRNAs by cytoplasmic injection of *in vitro* transcribed sgRNA and Cas9 mRNA into fertilized oocytes, as previously described^14^. Cutting was determined by extracting DNA from single blastocysts at day E4.5, amplifying the region around the Cas9 targeting site by PCR, and Sanger-sequencing the PCR product. In case of successful Cas9-mediated cleavage, insertions/deletions in one or both alleles are discernible as an altered pattern of chromatogram peaks (Fig. 1a). We defined as efficient any sgRNAs that cut at least 50% of blastocysts analyzed. Our final 3’ and 5’ sgRNAs cut 3/5 and 15/21 blastocysts, respectively (Fig.1b). The cut site for our final 5’ sgRNA (ID #6) was located 633 bp upstream of J_H_1, and the cut site for our 3’ sgRNA (ID #7) was located 108 bp downstream of J_H_4.

**Figure 1.**
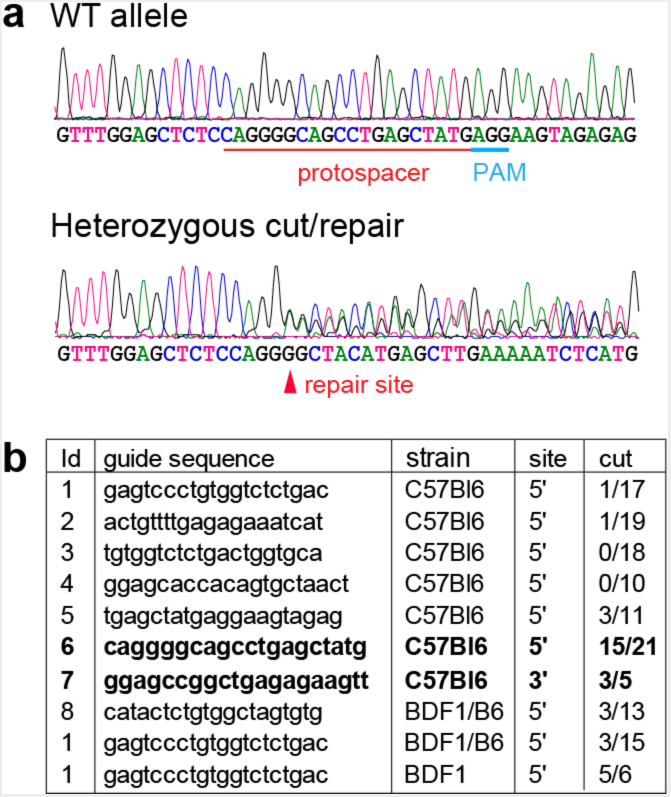
**Efficiency of sgRNAs flanking the mouse J_H_ region. (a)** Example sequences amplified from blastocysts 4 days after CRISPR/Cas9-mediated targeting by zygote injection. Top, WT (protospacer and PAM indicated) and bottom, successfully targeted blastocycts. Note the altered chromatogram peaks resulting from a monoallelic indel at the position indicated with an arrowhead (repair site). **(b)** List of tested sgRNA protospacer sequences, including mouse strain, location (5’ or 3’ of the J segments) and efficiency of cutting measured as in (a). The final sgRNAs used for generating knock-in mice are in bold font.

To build a monoallelic light/heavy chain Ig construct, we chose a B cell clone specific for the model antigen chicken gamma globulin (CGG; more specifically, the clone recognizes the constant region of IgY, the major component of CGG) that was efficiently recruited to germinal centers (GCs) upon CGG immunization in a polyclonal setting^15^. The targeting construct for the *Igh*^CGG^ allele consisted of an Ig V-region promoter followed by a pre-rearranged Igκ VJ segment, a human Igκ constant region (human Igκ^16^ was used for subsequent identification of cells bearing the transgenic receptor), a self-cleaving 2A peptide, and a pre-rearranged IgH VDJ segment (Fig. 2a). When targeted to the *Igh* J locus, this construct configuration results in the expression of both light- and heavy chain proteins from the same promoter, the heavy chain variable region being spliced onto the endogenous IgH constant region. The 2A peptide sequence allows for stoichiometric expression of the light and heavy immunoglobulin chains. To reduce focused AID targeting in the region encoding human Igκ and 2A peptide, we eliminated all AID hotspot motifs (RGYW) from this sequence by introduction of silent mutations (Fig. S1). This construct was cloned into a targeting vector containing 5’ and 3’ homology arms of 7.9 and 3.4 kb, respectively, which was previously used for generating *Igh* targeted insertions in C57BL6 embryonic stem cells^17^. As with the sgRNAs, the homology arms flank the endogenous J segments, removing this section of the *Igh* locus upon successful homology-directed repair and thus preventing further rearrangement of the targeted locus.

**Figure 2.**
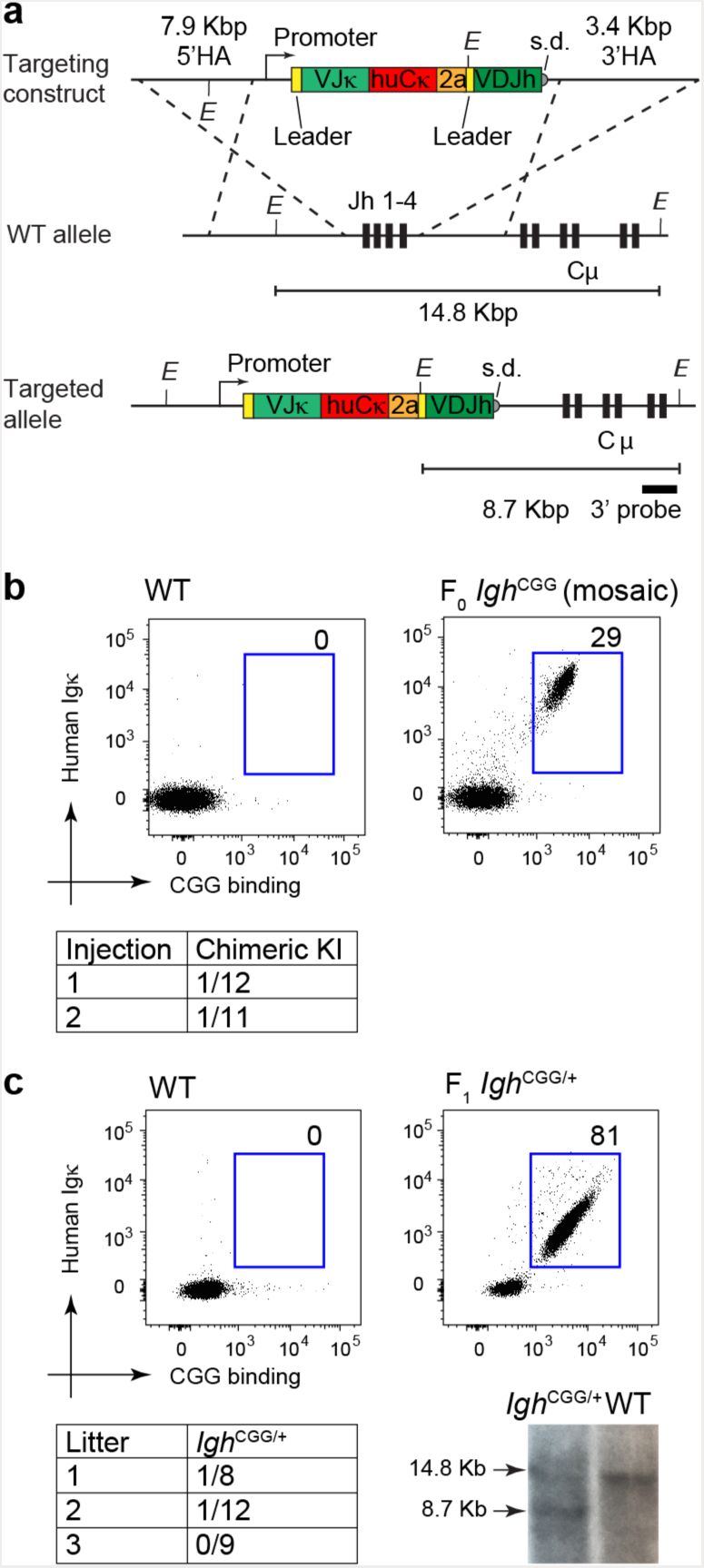
**Generation of a bicistronic VJC**κ**+VDJ_H_ anti-CGG knock-in mouse. (a)** Schematic representation of the bicistronic VJCκ+VDJ_H_ targeting cassette used to generate the *Igh*^CGG^ allele, before and after targeting to the *Igh* locus. The total insert size is 1.7 Kbp. The 3’ probe used for Southern blot is indicated. s.d., splice donor; E, EcoRV. Segments not drawn to scale. **(b)** Flow cytometry of blood samples, gated on B220^+^ cells, showing expression of human Igκ and CGG binding. One WT and one correctly targeted founder mouse are shown. Frequency of *Igh*^CGG^ positive founders in two rounds of injection is shown below (positive/total pups). **(c)** Germline transmission of *Igh*^CGG^ to F_1_ mice. One *Igh*^CGG^ founder female was mated to a C57BL6 male, and positive offspring in sequential litters were determined by flow cytometry as in (d). Frequency of *Igh*^CGG/+^ F_1_ pups (knock-in/total) in each litter is shown below. A Southern blot confirming integration into the *Igh* locus is also shown. An intact germline Igh locus yields a 14.8 kb fragment while integration into the locus yields a 8.7 kb fragment.

Cytoplasmic zygote injection of this construct along with Cas9mRNA, sgRNAs #6 and #7 (Fig. 1b) and an inhibitor of non-homologous end joining^18^ yielded 1/12 and 1/11 pups positive for human Igκ in two independent experiments. These F_0_ mice displayed a relatively low proportion of B cells carrying the engineered receptor (Fig. 2b), indicative of mosaicism resulting from targeting taking place after the first chromosome duplication. Upon breeding, human Igκ^+^ mice (*Igh*^CGG/+^) were born at sub-Mendelian ratios, and positive F_1_ mice carried a high proportion of B cells expressing the engineered receptor (Fig.2c), supporting the notion of mosaic targeting of the F_0_ mouse. We validated the integration of our construct into the *Igh* locus by Southern blotting of a heterozygous F_1_ mouse (Fig. 2c).

To determine whether presence of the *Igh*^CGG^ allele affected peripheral B cell populations, we analyzed the spleen of *Igh*^CGG/+^ mice for follicular (AA4.1^−^ CD23^+^ CD21^lo^ IgM^+^), marginal zone (MZ; AA4.1^−^ CD23^−^ CD21^hi^ IgM^hi^), and B1 (AA4.1^−^ CD23^−^ CD21^int^ IgM^int^) B cells, as previously described^19^. CGG-binding *Igh*^CGG/+^ B cells showed normal B cell subset distribution when compared to wild type (WT) mice. The single exception to this was a slightly decreased contribution of human Igκ+ CGG-binding cells to the MZ subset when compared to B cells from the same mouse that did not express the transgene (Fig. 3a,b). B cells expressing the pre-rearranged receptor did not co-express the endogenous mouse *Igk* gene, indicating appropriate exclusion of the endogenous *Igk* allele by the transgene (Fig. 3c). Mature follicular CGG-binding B cells showed slightly lower mean expression of surface IgM and a stronger reduction in mean expression of IgD when compared to WT polyclonal cells, although expression of both isotypes was still within the range observed in polyclonal cells (Fig. 3d). Lower surface Ig (sIg) expression was not due to overt failure of the 2A peptide to induce ribosome skipping and subsequent separation of light and heavy chains, since the full length VJCκ-VDJCμ protein (~95 kDa band) could not be detected by anti-IgM Western blot of naïve *Igh*^CGG/+^ B cells (Fig. 3e). While the reasons for lower sIg expression in these mice are unclear, sIg levels are likely to be a clonal property, in that different monoclonal B cell mice display different levels of sIg^6,20^. Therefore, we cannot determine whether the low sIg expression seen in *Igh*^CGG/+^ mice is a consequence of our knock-in strategy or of the specific CGG-reactive clone we used to generate this strain. Future monoclonal strains generated using this strategy with different Ig sequences will help resolve this issue.

**Figure 3.**
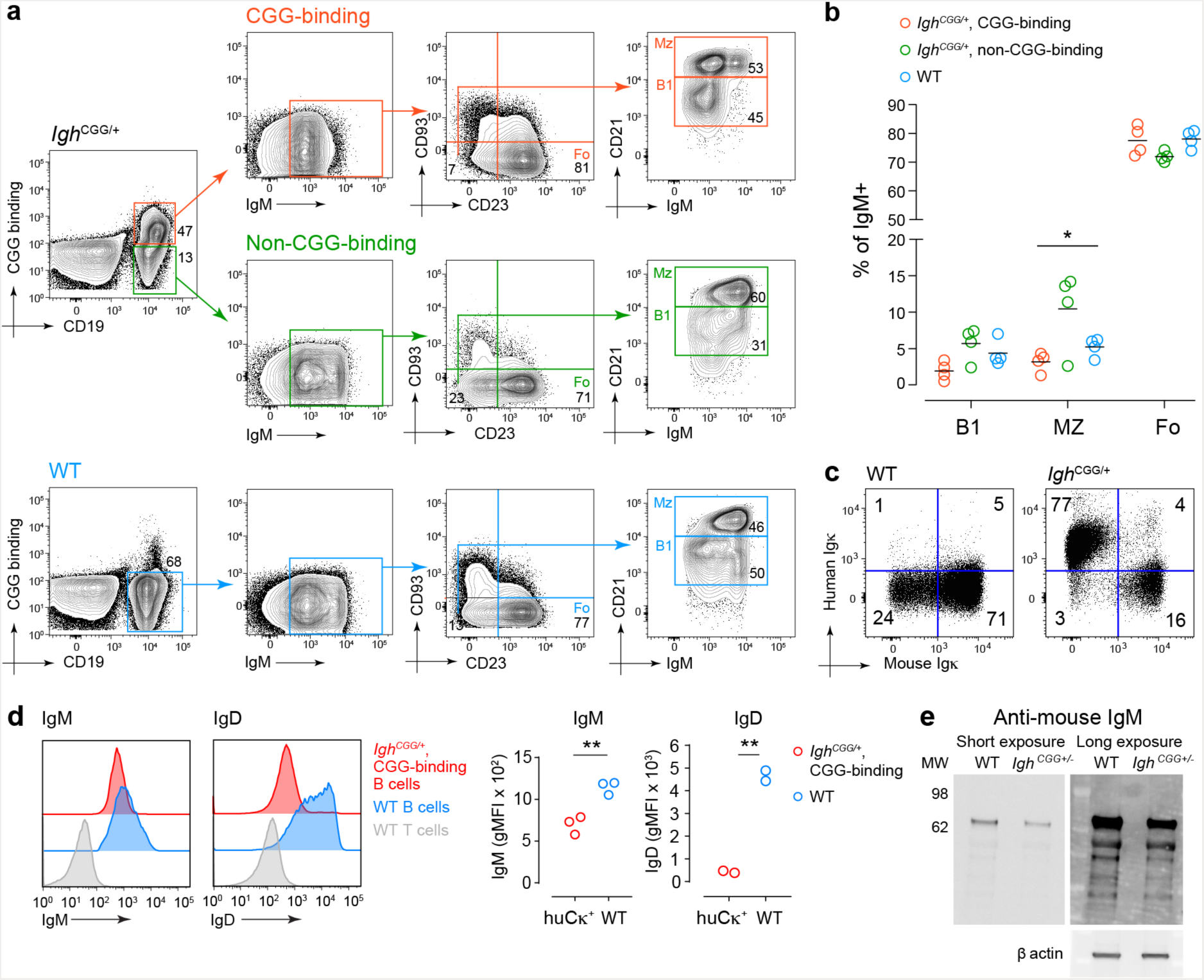
**B cell subsets and surface Ig expression in *Igh*^CGG/+^ mice. (a)** Splenocytes from 6-week-old *Igh*^CGG/+^ or WT C57BL6 mice were stained as indicated. For *Igh*^CGG/+^ mice, CD19^+^ cells were analyzed as either CGG-binding or non-binding (respectively marked as I. or II., the latter representing B cells that escaped the transgene). Cells were further gated as IgM^+^ and analyzed for CD23 and CD93 expression. Cells negative for CD93 and CD23 (LL, Lower Left quadrant)) were further analyzed for IgM and CD21. From this plot Marginal Zone cells (MZ: CD93^−^ CD23^−^ CD21^hi^ IgM^hi^) and B1 cells (B1: CD93^−^ CD23^−^ CD21^int^ IgM^int^) were identified. Cells negative for CD93 and positive for CD23 (Lower Right, LR quadrant) were analyzed for IgM and CD21 with the identification of follicular B cells (Fo: AA4.1^−^ CD23^+^ CD21^lo^ IgM^+^). WT mice were analyzed identically on the CD19^+^/CGG^neg^ population. **(b)** Summary of data as in (a) for 4 mice of each genotype. MZ, B1 and Fo B cells are shown as percentage of all IgM^+^ cells for WT mice or as percentage of IgM+ cells of that class (CGG-binding or non-binding) *Igh*^CGG/+^ mice. *p = 0.039 by unpaired t-test. **(c)** Flow cytometry of blood samples showing allelic exclusion of endogenous mouse Igκ in *Igh*^CGG/+^. Gated on B220^+^. One plot representative of three mice is shown. **(d)** Surface IgM and IgD expression on WT and *Igh^CGG/+^* follicular B cells. Gated on CD19^+^/huCκ^+^/CD93^neg^/CD23^+^ cells for *Igh^CGG/+^* mice or on CD19^+^/CD93^neg^/CD23^+^ cells for WT. WT T cells are shown as a negative control. Data from independent experiments are shown on the right. Each symbol represents one mouse. ** p = 0.0046 (IgM) or p = 0.0030 (IgD) by unpaired t test. gMFI, geometric mean fluorescence. **(e)** Western blot for IgM in naïve B cells from *Igh*^CGG/+^ mice. Anti-IgM signal for WT and anti CGG B cells is shown at two different exposures. The relevant band is indicated at around 70 KDa. The β actin loading control is shown below. Note the absence of a ~95 kb band, the expected size of the uncleaved Vκ/V_H_ construct. Representative of two experiments.

We next sought to determine whether *Igh*^CGG/+^ B cells were fully functional in spite of their lower expression of sIg. We first cultured splenic B cells *in vitro* with LPS and IL-4 to induce isotype switching to IgG_1_. *Igh*^CGG/+^ B cells switched to IgG_1_ at WT levels, although surface expression of this isotype was again slightly lower than WT (Fig. 4a). We then adoptively transferred *Igh*^CGG/+^ B cells into a CD45.1 congenic strain, which we immunized with CGG precipitated in alum. *Igh*^CGG/+^ B cells were able to efficiently access germinal centers (GCs) and class switch to IgG_1_, indicating that *Igh*^CGG^ B cells are capable of proper progression through all steps of B cell activation. Again, surface expression of IgG_1_ on CGG-binding GC B cells was slightly lower than that of their WT counterparts (Fig. 4b). Thus, despite lower expression of sIg, *Igh*^CGG/+^ cells are fully competent to engage in GC reactions and undergo CSR *in vivo*.

**Figure 4.**
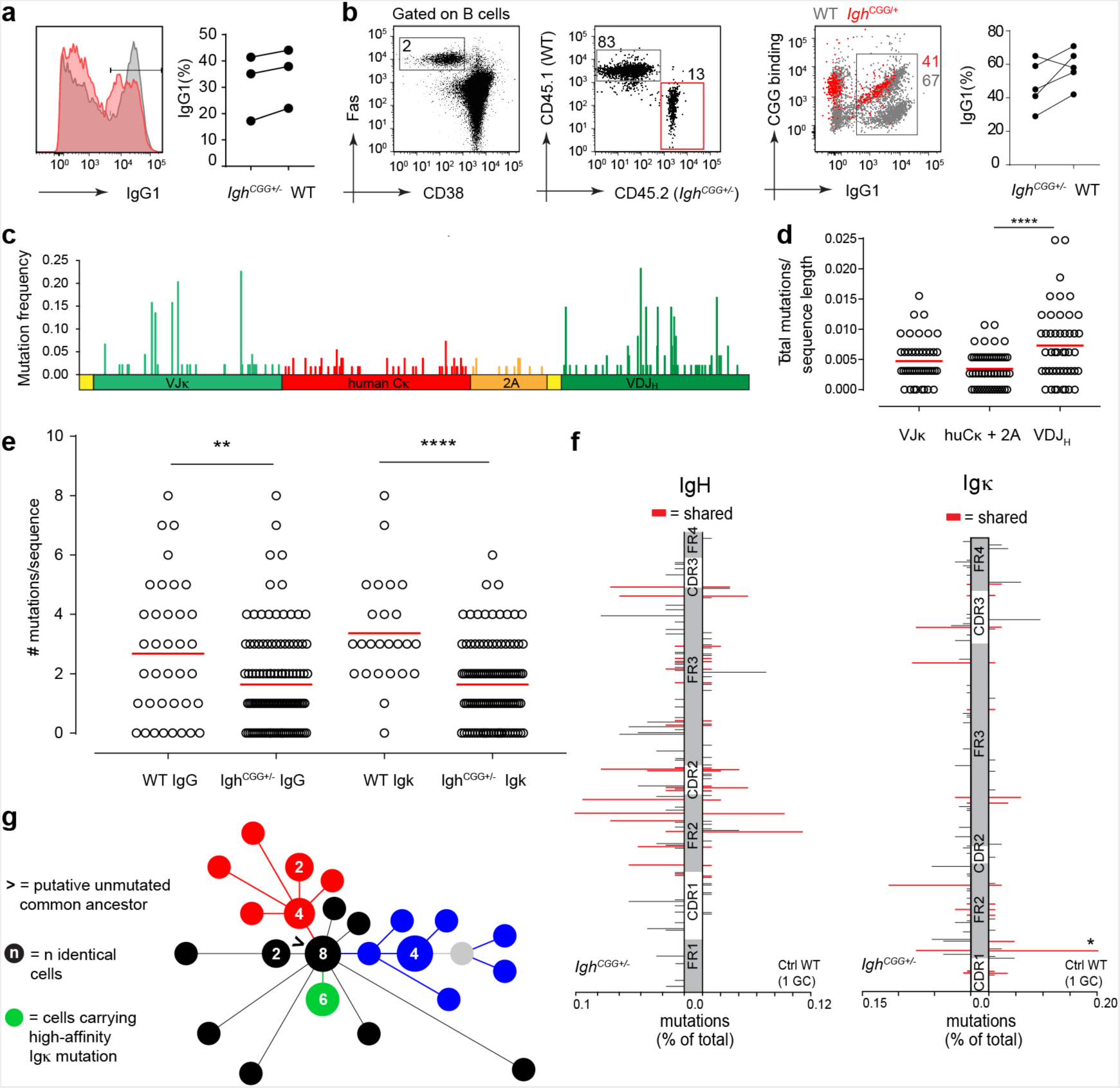
**Class-switch recombination, somatic hypermutation, and antigen-driven selection in *Igh*^CGG/+^ B cells. (a)** In vitro isotype switching of *Igh^CGG/+^* B cells. Purified *Igh*^CGG/+^ (CD45.2/2) and WT (CD45.1/1) B cells were co-cultured *in vitro* with LPS and IL-4 to induce isotype switching to IgG_1_. Histogram shows IgG_1_ staining by flow cytometry on day 3 of culture. Data from 3 independent experiments are quantified in the graph; proportions of IgG_1_^+^ cells in *Igh*^CGG/+^ and WT B cells from the same experiment are connected by a line. P (ns)= 0.74 by unpaired t test. **(b)** *Igh*^CGG/+^ B cells (CD45.2/2) were adoptively transferred into WT (CD45.1/1) recipients which were immunized one day later with CGG in alum. CSR to IgG^1^ was determined in GC B cells (CD19^+^TCRβ^−^Fas^+^CD38^lo^) from donor (CD45.2/2) and recipient (CD45.1/1). Graph shows percentage of IgG_1_^+^ GC B cells in 5 mice from two independent experiments. Proportions of switched cells in the same mouse are connected by a line. P (ns)= 0.23 by unpaired t test. **(c-f)** GC B cells from adoptive transfer experiments similar to in (d) were single-sorted and the entire *Igh*^CGG/+^ allele was PCR-amplified and sequenced. **(c)** Nucleotide mutation frequencies along the *Igh*^CGG/+^ locus, calculated as the fraction of times a particular nucleotide was mutated from the original sequence. **(d)** Total mutation frequency per region (normalized to sequence length). Each symbol represents one cell. p<0.0001 for VDJ_H_ compared to huCκ+2A and p (ns)= 0.0602 for VJk compared to huCκ+2A by unpaired t test. **(e)** Number of nucleotide mutations in VDJ_H_ and VJκ in *Igh*^CGG/+^ GC B cells compared to WT GC B cells carrying the same rearrangement selected by CGG immunization, reported previously (^15^; only data from LN#2 GC2 in Fig.4 from ^15^ is analyzed). Data from two transfers of *Igh^CGG/+^* B cells are pooled and compared to the WT: IgG **p=0.0026 and Igκ ****p<0.0001, by unpaired t-test **(f)** Comparison of nucleotide mutation patterns between *Igh*^CGG/+^ B cells and WT GC B cells carrying the same rearrangement. Mutation patterns are shown for FR1 to FR4 regions of IgH and CDR1 to FR4 regions of Igκ. Shared nucleotide mutation positions are shown in red. *= C119>G mutation, which confers an ~8-fold gain in affinity. Data from two transfers of *Igh^CGG/+^* B cells are pooled and compared to the WT **(g)** Reconstructed pyhologeny of *Igh*^CGG/+^ GC B cell VJ Igκ sequences from multiple GCs in one lymph node. Clades of expanded clones are indicated in red, green and blue. Cells containing the high affinity C119>G mutation are indicated in green.

To assess the ability of *Igh*^CGG/+^ B cells to undergo somatic hypermutation and affinity maturation, we analyzed the mutation patterns of single GC B cells sorted from experiments analogous to those detailed in Fig. 4b. *Igh*^CGG/+^ B cells accumulated substantial SHM across the entire engineered locus (Fig. 4c-e). Removal of AID hotspots from human *Igk* and 2A peptide led to a slight reduction in SHM in these regions (Fig. 4c,e), compatible with the partial requirement for the “hotspot” motif in AID targeting^21^. The accumulation of mutations at similar positions (tall peaks in Fig. 4c) in VJκ and VDJ_H_, but not in Cκ or 2A suggested that antigen-based selection was active on this allele. To verify this, and to determine whether SHM burdens in *Igh*^CGG/+^ B cells were comparable to those of WT B cells, we took advantage of the fact that we had previously characterized the SHM and evolution of this particular anti-CGG clone in two separate GCs *in vivo*^15^. We therefore have a previous record of the maturation of this very clone under WT conditions, with which we could compare the evolution of our transgenic B cells. To avoid unfair comparisons imposed by the presence of a strong selective bottleneck, we chose a GC in which this clone was expanded in several parallel sub-lineages, rather than by a single selective sweep (Fig.4 “LN#2 GC#2”^15^). We then compared the SHM pattern of cells from this GC to that of *Igh*^CGG/+^ GC B cells pooled from two whole lymph nodes. This analysis showed that SHM in *Igh*^CGG/+^ B cells reached a substantial fraction of the levels seen in this particular GC at the same time point (49% for *Igk* VJ and 61% for *Igh* VDJ, Fig. 4e). Importantly, comparison of WT and *Igh*^CGG/+^ B cells showed a remarkable coincidence in the most frequently mutated positions (red bars in Fig. 4f), including accumulation of an Igκ mutation (C119>G) showed to confer an 8-fold increase in affinity to this clone and that was strongly selected in WT GCs^15^. By reconstructing phylogenic trees of VJκ, we could identify three separate clusters of expanded sub-clones (suggestive of homogenizing selection events occurring in three separate GCs), one of which included the known high-affinity light chain mutation in position 119 (Fig. 4g). Thus, *Igh*^CGG/+^ B cells are capable of somatic hypermutation and antigen-driven selection to an extent similar to that of WT B cells bearing the same Ig rearrangements.

Here, we report the development of a hybrid method that occupies an intermediate position between randomly inserted Ig transgenes^1^ and full two-chain Ig knock-ins. As with randomly inserted transgenes, our monoallelic Ig mice can be generated within 2-3 months, and do not require the transmission of two alleles, greatly expediting mouse generation and subsequent breeding to other genetically modified strains. Whereas sIg expression for this particular anti-GCC BCR is lower than the average sIgM and sIgD expression by polyclonal B cells from WT mice, SHM, CSR, and affinity-based selection—arguably the key reasons to use mice with Ig knock-in alleles rather than randomly inserted Ig transgenes—are largely preserved. We therefore expect that monoallelic Ig knock-in mice could greatly facilitate studies that require a generation of a large panel of monoclonal B cell receptor mice, including studies of the origin and evolution of broadly neutralizing antibodies in HIV ^22^ or of pathogenic autoantibodies in autoimmune disease ^23^.

## MATERIALS AND METHODS

### Design and cloning of the targeting construct

To generate the targeting construct we first synthesized a shuttle vector containing a ~500 bp IgH-V promoter, a V_H_ leader sequence followed by an XhoI site, a human Cκ chain and the porcine teschovirus-1 2A peptide (both re-coded to eliminate AID hotspots), a second V_H_ leader sequence followed by an AflII site, and part of the J_H_4 intron including 50 bp starting from the splice donor (Fig. S1). To avoid repetitive sequences that would prevent gene synthesis, one of the V_H_ leaders was recoded. This fragment was cloned into Zero Topo blunt (Thermo Fisher, #451245). Subsequently the VJκ of the anti-CGG antibody^15^ was cloned into the XhoI site and the VDJ_H_ of the same antibody was cloned into the AflII site. This full construct was cloned into a previously designed vector containing 3’ and 5’ homology arms for insertion into the Igh locus^24^. As the original vector with the IgH locus homology arms^24^ was used for generating ES cell lines with IgH insertions, it contained a Diphtheria Toxin A (DTA) selection cassette, which we removed by subcloning the full targeting insert (5’ homology arm – Ig transgene – 3’ homology arm) into pBluescript KS. For injection the final plasmid was purified using an endotoxin free maxiprep kit (Qiagen, #12362).

### sgRNA generation

For making each sgRNA a gBlock with a built-in T7 priming sequence and the guide/scaffold sequence was synthesized using the following sequence: CGCTGTTAATACGACTCACTATAGGGn_(20)_ GTTTTAGAGCTAGAAATAGCAAGTTAAAATAAGGCTAGTCCGTTATCAACTTGAAAAAGTGGCACCGAGTCGGTGCTTTT where n_(20)_ represents the location of the 20 bp CRISPR RNA sequence. The lyophilized gblock was reconstituted in 10 µl of PCR-grade water to make 20 ng/µl stock. RNA was synthesized *in vitro* from the gblock using MEGAshortscript T7 kit (AM1354M, LifeTech), using 8 µl of the reconstituted gblock as a template. To clean up the RNA for synthesis, we used Agencourt RNAClean XP beads (A63987, Agencourt). We added 50 µl of beads for the synthesized RNA and incubated for 10 minutes. Using a 95 w plate magnet, the beads were washed 3 times in 80% EtOH. After the final EtOH wash beads were briefly left to dry and resupended in HyClone Molecular Biology-Grade Water (SH30538.02, GE Healthcare Life sciences). Beads were removed from the solution after a 5 min incubation. The RNA was stored at -80 until use.

### Testing sgRNA cutting efficiency in blastocysts

To test the effiency of our sgRNA, they were injected individually into blastocysts (as detailed below). Blastocysts were cultured until E4.5. and collected into tubes with 5µl QuickExtract buffer (Epicentre, QE09050). To obtain PCR ready genomic DNA, we incubated the tubes at 65^o^C for 6 min, followed by a quick vortex and a 2 min incubation at 98^o^C. The entire 5 µl was used as a PCR template. For the 1 kb sequence spanning the 5’ cutting site we used the following primers: forward 5’agagatactgcttcatcaca3’ and reverse 5’gggacctgcacctatcctgtcc3’. For the 1 kb sequence spanning the 3’ cut site we used the following primers: forward 5’ataggttatgagagagcctcac3’ and reverse 5’ctgacagttgatggtgacaatt3’. We performed a PCR with Taq polymerase (M0273, New England Biolabs); 95^o^C for 30 s, 30 cycles of 95^o^C for 30 s, 58^o^C for 60 s and 68^o^C for 30 s, and a final extension time of 68^o^C for 5 min. PCR products were Sanger sequenced using their respective forward primers.

### Zygote Injections

Zygote injections were performed at the Genetically Engineered Models (GEM) Center of the Whitehead Institute for Biomedical Research. All procedures were performed according to NIH guidelines and approved by the Committee on Animal Care at MIT. Super-ovulated 8-10 wk old female B6D2F2 mice ((7.5 IU of pregnant mare serum gonadotropin (367222-1000IU, EMD Millipore), followed 46-48h later by 7.5 IU chorionic gonadotropin (80051-032, VWR)) were mated to stud males and fertilized pronuclear stage embryos (zygotes) were collected approximately 20h post hCG. Cytoplasmic injections were performed using a Piezo actuator (PMM-150FU, Prime Tech) and a flat tip microinjection pipette with an internal diameter of 8 µm (Origio Inc, Charlottesville, VA). The injection mix was prepared immediately prior to the procedure and included the following components at the final concentrations indicated: 100 ng/µL Cas 9 mRNA (CAS9MRNA-1EA, Sigma Aldrich), 50 ng/µL sgRNA, 200 ng/µL of our IgH targeting plasmid and 1 mM SCR-7 (M60082-2s, Excess Biosciences) that inhibits Non-Homologous End Joining^18^ following the procedure published previously^25^. Immediately after the completion of the injection, approximately 20 zygotes were transferred into the oviducts of pseudopregnant ICR females (CD-1, Charles River) at 0.5 dpc.

### Western blotting

B cells were purified from spleen with Miltenyi CD45R isolation beads (130-049-501, Miltenyi) with 0.5 x10^6 cells used for every well. The cells were incubated in RIPA buffer for 2 minutes at RT. Loading buffer (Invitrogen, NP0007) and sample buffer (Invitrogen, NP0009) was added to the samples. The samples were sonicated for 5 minutes and then spun down for 2 minutes. The supernatant was incubated at 90ºC for 5 minutes. The samples were run on a 4-12% bis-tris NuPage gel (Thermo Fisher, NP0322) with 7 µl prestained SeeBlue Plus2 (Invitrogen, LC5925). The gel was run in MOPS running buffer at 120v for 1.5 h then blotted onto a PVDF membrane (Millipore Immobilion PVDF membrane, IPVH20200) in transfer buffer at 4ºC for 1 h at 300 mA. After transfer, the membrane was incubated in blocking buffer at RT for 2 h. Goat anti mouse IgM (1:6,000) ^26^ was added to the blocking buffer and incubated for 1 h at RT. The membrane was rinsed three times with PBS 0.05% Tween20 (Millipore Sigma, P9416). After drying, 5-6 ml of Hyglo Quick spray (Denville scientific Inc, E2400). Chemiluminescence was detected after a 30 seconds or 20 minutes incubation. For the loading controls, the membrane was stripped and restained with 1:15,000 anti beta actin (GeneTex, GTX109639).

### Southern blotting

A southern blot was performed as described previously ^17^. Genomic DNA was digested with EcoRV.

### Cell transfers and immunizations

Adoptive transfer of anti-CGG B cells was carried out either by transferring blood (Fig.2c) or purified B cells (Fig.2b) from donor to recipient. For B cells transferred in blood, 100 µl of blood was collected from a donor mouse and directly injected intravenously into the recipient. 100 µl of blood corresponds to approximately 4 x 10^5^ B cells. For transfer of purified B cells, we isolated B cells from total splenocyte preparations by negative selection using anti-CD43-coupled magnetic beads (130-090-862, Miltenyi Biotec). Untouched B cells were purified according to the manufacturer’s protocol and 5 x 10^5^ B cells were transferred into each recipient. 24 h following cell transfers, mice were immunized in each footpad with CGG (10 µg) precipitated in 1/3 volume Imject Alum (77161, ThermoFischer scientific). To distinguish our transferred B cells, we used CD45.1 congenic mice as recipients (Jackson Laboratories, strain 002014).

### Sample processing for flow cytometry and cell sorting

LNs were placed in microcentrifuge tubes containing 100 µl of PBS supplemented with 0.5% BSA and 1mM EDTA (PBE), macerated using disposable micropestles (AXYPES-15-B-SI,Axygen), and further dissociated into single-cell suspensions by gentle vortexing. 100 µl of 2X antibody stain (antibodies to CD38, IgD, FAS, B220, CD4 supplemented with Fc block) was added to the cell suspension, which was incubated on ice for 30 min. Single cells were sorted into 96-well plates, as described below, using a FACS Aria II cell sorter.

### Antibodies for flow cytometry analysis

See supplementary table 1 for detailed information about antibodies used for phenotyping CGG B cell mice, adoptive transfer and isotype switching experiments. *Single-cell Igh and Igk PCR and sequence analysis* Sorted single cells processed and analyzed essentially as described previously^15^. Single cell variable regions were amplified by semi nested PCR using 5’ATGGGtTGGTCCTGCATTATACTGT3’ (fw L-VH recoded) with 5’ggaaggtgtgcacaccgctggac3’ (rev IgG1) and 5’AGGGGGCTCTCGCAGGAGACGAGG3’ (rev IgM) for the first PCR and fw and 5’GCTCAGGGAAATARCCCTTGAC3’ (IgG1 internal rev) for the second PCR. PCRs were performed with Taq polymerase (NEB, M0273). Products were amplified with the following program 94^o^C for 2 min, 29 rounds of 94^o^C for 30 s, 56^o^C for 30 s and 72^o^C for 55 s, followed by a final extension of 72^o^C for 10 min. Products were sent for sequencing with the following primers: 5’GAATGTACACCGGTTGCAGTTGCTA3’ to obtain huCk and 2A, 5’ATGGGtTGGTCcTGcATtATaCTgT3’ to obtain VJ and 5’GCTCAGGGAAATARCCCTTGAC3’ (IgG1 internal rev) to obtain VDJ. Analysis of flow cytometry data for presentation was carried out using FlowJo software, v. 10.

### Isotype switching assay

*In vitro* isotype switching was essentially performed as previously described ^27^. Untouched B cells were isolated using Miltenyi separation beads and columns and cultured at 3 x10^5^ cells/ml in 24 well tissue culture coated plates. The cells were cultured in complete medium: RPMI-1640 medium plus glutamine (10-040-CV, Corning cellgro) containing 100 U/mL penicillin and 100 µg/mL streptomycin (P11-010, PAA Laboratories), 10% fetal calf serum (FCS; heat inactivated, SH30910.03, GE Healthcare Life Sciences), and 5 x 10^-5^ M 2-mercaptoethanol (ME) (M7552,Sigma). 20 ng/ml recombinant IL-4 (574302, Biolegend) and 40 ng/ml LPS (Escherichia coli LPS serotype 055:B5 L2880, Sigma). The cells were stained for IgG_1_ on day 4.

### Statistical analysis

Differences in means for two-sample comparisons were evaluated using the two-tailed Student’s t-test.

## Acknowledgements

We thank Yotam Bar On and the gene targeting resource center (Rockefeller University) for technical advice and assistance. This work was funded by NIAID grant R01AI119006 and by the Searle Scholars Program (both to G.D.V.), The Research Council of Norway, pr.no 239757, co-funded by the European Union’s Seventh Framework Programme for research, technological development and demonstration under Marie Curie grant agreement no 608695 (to J.T.J) and CAPES fellowship/PDSE/process number 88881.132337/2016-01 (to C.B.C).

---

**Supplementary Figure 1.**
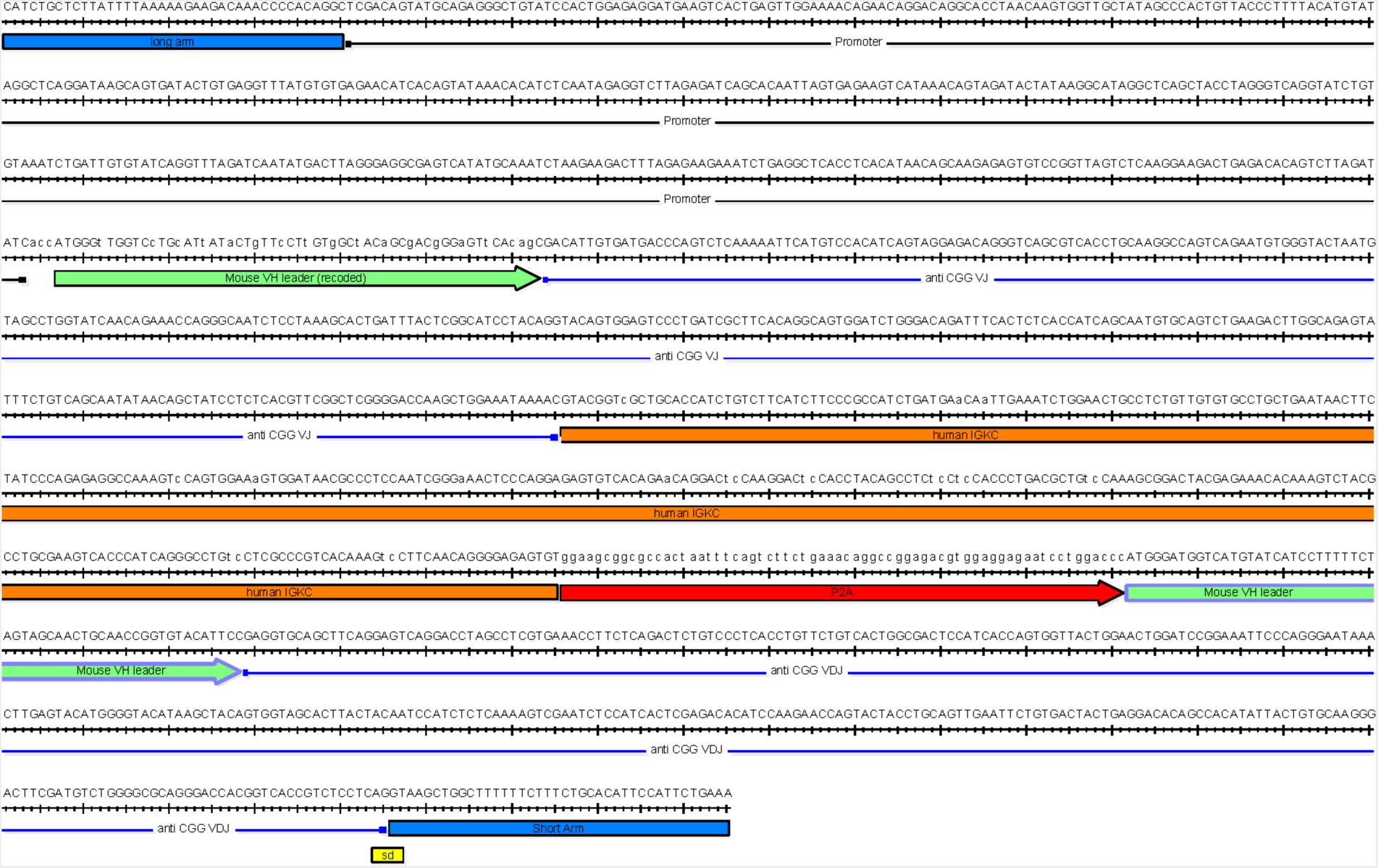
Sequence of the *Igh*^CGG^ insertion. Terminal ends of the 5’ (long) and 3’ (short) arm of homology are shown in blue. The recoded leader for the anti CGG VJ is shown in green with altered bases indicated in lower case letters. The coding sequence for human Igκ is shown in orange, with silent mutations used to eliminate AID hotspots indicated as lower-case letters. The splice donor (sd) site is indicated in the 3’ untranslated region of anti CGG VDJ.

**Supplementary Table 1.**
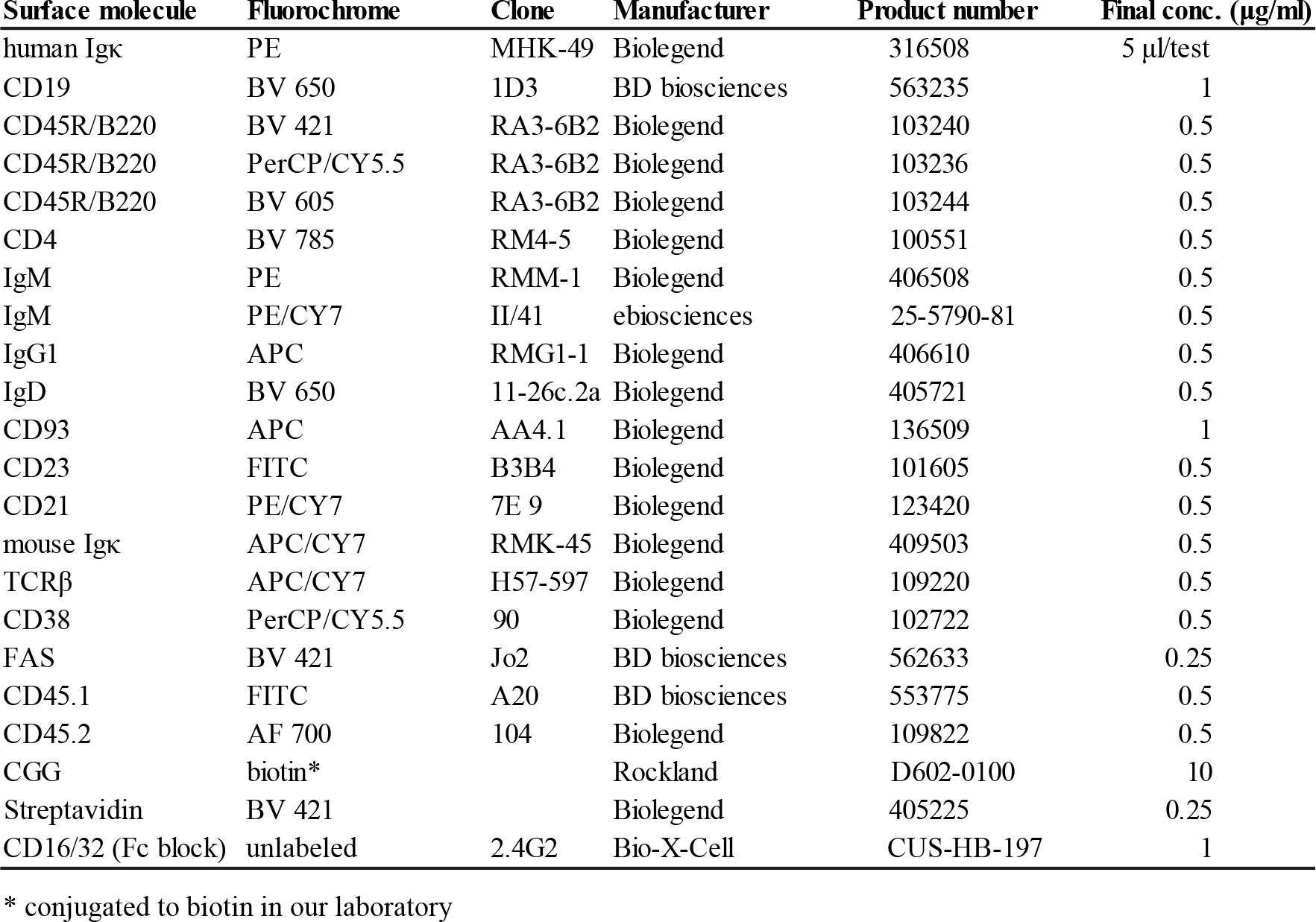
Reagents used for flow cytometry.

| Surface molecule | Fluorochrome | Clone | Manufacturer | Product number | Final conc. (µg/ml) |
| --- | --- | --- | --- | --- | --- |
| human Igκ | PE | MHK-49 | Biolegend | 316508 | 5 µl/test |
| CD19 | BV 650 | 1D3 | BD biosciences | 563235 | 1 |
| CD45R/B220 | BV 421 | RA3-6B2 | Biolegend | 103240 | 0.5 |
| CD45R/B220 | PerCP/CY5.5 | RA3-6B2 | Biolegend | 103236 | 0.5 |
| CD45R/B220 | BV 605 | RA3-6B2 | Biolegend | 103244 | 0.5 |
| CD4 | BV 785 | RM4-5 | Biolegend | 100551 | 0.5 |
| IgM | PE | RMM-1 | Biolegend | 406508 | 0.5 |
| IgM | PE/CY7 | II/41 | ebiosciences | 25-5790-81 | 0.5 |
| IgG1 | APC | RMG1-1 | Biolegend | 406610 | 0.5 |
| IgD | BV 650 | 11-26c.2a | Biolegend | 405721 | 0.5 |
| CD93 | APC | AA4.1 | Biolegend | 136509 | 1 |
| CD23 | FITC | B3B4 | Biolegend | 101605 | 0.5 |
| CD21 | PE/CY7 | 7E 9 | Biolegend | 123420 | 0.5 |
| mouse Igκ | APC/CY7 | RMK-45 | Biolegend | 409503 | 0.5 |
| TCRβ | APC/CY7 | H57-597 | Biolegend | 109220 | 0.5 |
| CD38 | PerCP/CY5.5 | 90 | Biolegend | 102722 | 0.5 |
| FAS | BV 421 | Jo2 | BD biosciences | 562633 | 0.25 |
| CD45.1 | FITC | A20 | BD biosciences | 553775 | 0.5 |
| CD45.2 | AF 700 | 104 | Biolegend | 109822 | 0.5 |
| CGG | biotin* |  | Rockland | D602-0100 | 10 |
| Streptavidin | BV 421 |  | Biolegend | 405225 | 0.25 |
| CD16/32 (Fc block) | unlabeled | 2.4G2 | Bio-X-Cell | CUS-HB-197 | 1 |

\* conjugated to biotin in our laboratory

* conjugated to biotin in our laboratory

